# Mutual Potentiation of Plant Immunity by Cell-surface and Intracellular Receptors

**DOI:** 10.1101/2020.04.10.034173

**Authors:** Bruno Pok Man Ngou, Hee-Kyung Ahn, Pingtao Ding, Jonathan DG Jones

## Abstract

The plant immune system involves cell-surface receptors that detect intercellular pathogen-derived molecules, and intracellular receptors that activate immunity upon detection of pathogen-secreted effectors that act inside the plant cell. Surface receptor-mediated immunity has been extensively studied but in authentic interactions between plants and microbial pathogens, its presence impedes study of intracellular receptor-mediated immunity alone. How these two immune pathways interact is poorly understood. Here, we reveal mutual potentiation between these two recognition-dependent defense pathways. Recognition by surface receptors activates multiple protein kinases and NADPH oxidases, whereas intracellular receptors primarily elevate abundance of these proteins. Reciprocally, the intracellular receptor-dependent hypersensitive cell death response is strongly enhanced by activation of surface receptors. Activation of either immune system alone is insufficient to provide effective resistance against *Pseudomonas syringae*. Thus, immune pathways activated by cell-surface and intracellular receptors mutually potentiate to activate strong defense that thwarts pathogens. By studying the activation of intracellular receptors in the absence of surface receptor-mediated immunity, we have dissected the relationship between the two distinct immune systems. These findings reshape our understanding of plant immunity and have broad implications for crop improvement.

## Main Text

In plants, innate immunity involves both cell-surface and intracellular receptors that upon recognition of pathogen-derived molecules, initiate immune responses^1^. Plant cell-surface pattern-recognition receptors (PRRs) recognize pathogen-associated molecular patterns (PAMPs) and signal via both plasma-membrane-associated co-receptor kinases, such as BAK1, and receptor-like cytoplasmic kinases (RLCK), such as BIK1^2^. Ligand-dependent association between PRRs and these protein kinases activates multiple cellular changes including calcium influx, production of reactive oxygen species (ROS) via the activation of plant NADPH oxidases encoded by respiratory burst oxidase homolog (Rboh) genes, activation of mitogen-activated protein kinases (MAPKs) and induction of defense genes^2–4^.

Intracellular nucleotide-binding-leucine-rich-repeat (NLR) receptors activate immune responses upon recognition of effectors secreted into host cells by pathogens, but in contrast to PRR responses, NLR-mediated responses are poorly defined. Plant NLRs are classified into two main classes based on either an N-terminal coiled-coil (CC) domain in CC-NLRs, or an N-terminal Toll/Interleukin receptor/Resistance protein (TIR) domain in TIR-NLRs^5,6^. Upon activation, the CC-NLR ZAR1 forms inflammasome-like complexes^7^ and associates with membranes. TIR-NLR defense activation requires their NADase activity^8,9^. PRR-mediated signaling is usually referred to as pattern-triggered immunity (PTI), and NLR-mediated signaling as effector-triggered immunity (ETI), though other terms with similar meanings have been proposed^10,11^. Despite recent progress in understanding immune receptor activation, our understanding of how PTI and ETI co-function to protect plants from pathogens is incomplete.

### ROS production induced by PTI is enhanced by ETI

To study ETI in the absence of PTI, we generated a transgenic Arabidopsis (*Arabidopsis thaliana*) line with estradiol-inducible expression of bacterial effector AvrRps4 recognized by intracellular TIR-NLRs RRS1 and RPS4. Estradiol application activates ETI^AvrRps4^. Pre-activation of ETI^AvrRps4^ elevates plant resistance against virulent bacteria *Pseudomonas syringae* pv. *tomato* (*Pst*) DC3000^12^. We tested if this enhanced resistance is due to potentiation of PTI responses by ETI^AvrRps4^. To investigate the effects of ETI on PTI, we pre-activated ETI^AvrRps4^ with estradiol and measured ROS production induced by a bacterial PAMP, flagellin-derived peptide flg22 (Extended Data Fig 1A). Pre-activation of ETI^AvrRps4^ leads to elevated ROS production during PTI, but induction of ETI alone does not activate ROS production (Extended Data Fig 1B and C). These data indicate that ETI elevates the strength of PTI responses.

During bacterial infection, activation of PTI precedes pathogen effector delivery into the host cell. To mimic this, we treated plants with flg22, or estradiol, or flg22 + estradiol, to activate PTI, or ETI^AvrRps4^ or “PTI + ETI^AvrRps4^”, respectively (Extended Data Fig 2A). We measured ROS production over 24 hours (h); “PTI + ETI^AvrRps4^” shows a much stronger ROS accumulation compared to PTI activation alone, particularly during the third phase (P-III) of the burst. Again, ETI^AvrRps4^ alone does not trigger ROS production (Fig 1A, B and Extended Data Fig 2B-G). We investigated if ETI mediated by CC-NLRs also potentiates PTI. The Arabidopsis CC-NLR RPS2 confers recognition of the bacterial effector AvrRpt2^13^. Using an estradiol-inducible AvrRpt2-expressing Arabidopsis line, we showed that ETI^AvrRpt2^ can also potentiate flg22-induced ROS burst, especially during P-II (Extended Data Fig 2H-O). Thus, ETI co-activation by TIR- or CC-NLRs enhances ROS production induced by PTI.

**Fig. 1.**
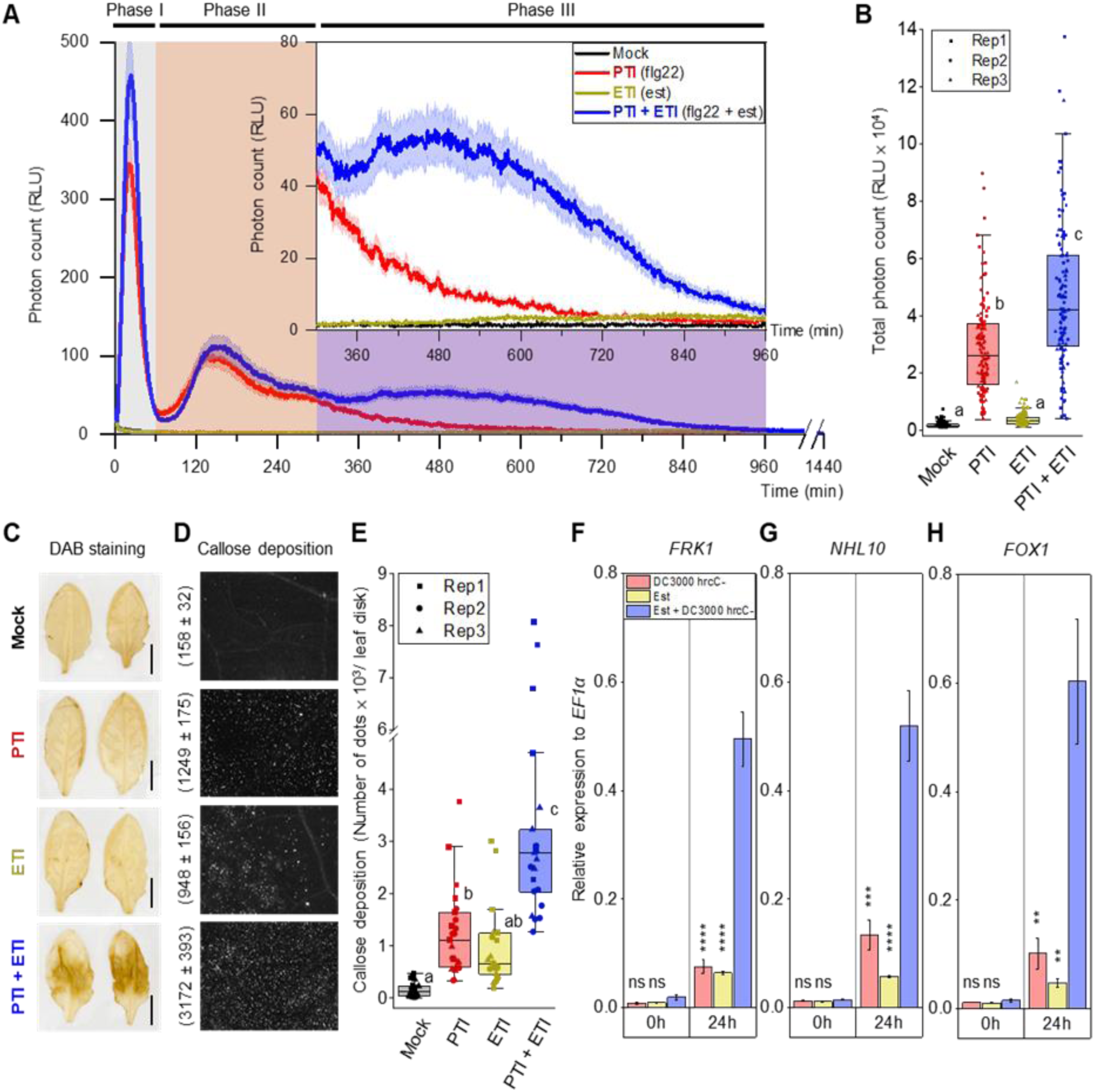
Potentiation of PTI-induced physiological changes during ETI^AvrRps4^. (A) “PTI + ETI^AvrRps4^” leads to prolonged ROS production from 300-960 minutes (5-16 h, phase III). Shaded curve represents standard error (S.E.). (B) Total ROS production in “PTI + ETI^AvrRps4^”-treated leaves is significantly higher than PTI-treated leaves. (C) “PTI + ETI^AvrRps4^” leads to higher hydrogen peroxide accumulation than PTI or ETI^AvrRps4^ alone. Scale bars represent 0.5 cm. (D) “PTI + ETI^AvrRps4^” leads to stronger callose deposition than PTI or ETI^AvrRps4^ alone. Numbers represent the mean and S.E. of a total of 23 leaves from each treatment. (F) Callose deposition in “PTI + ETI^AvrRps4^”-treated leaves is significantly higher than PTI- or ETI^AvrRps4^-treated leaves. (B, F) Data points from 3 biological replicates were analyzed with one-way ANOVA followed by Tukey’s HSD test. Data points with different letters indicate significant differences of P < 0.01. “PTI + ETI^AvrRps4^” leads to a stronger (F) *FRK1*, (G) *NHL10*, (H) *FOX1* (*AT1G26380*) transcript accumulation compared to PTI or ETI^AvrRps4^ alone. The average of data points from 3 biological replicates were plotted onto the graphs, with ±S.E. for error bars. Student’s t-test was used to analyze significance differences between PTI + ETI^AvrRps4^ and PTI or ETI^AvrRps4^. (ns, not significant; *; P ≤ 0.05; **, P ≤ 0.01; ***, P ≤ 0.005; ****, P ≤ 0.001).

As “PTI + ETI” leads to a stronger ROS burst than PTI alone, we monitored ROS accumulation in leaves by assessing the levels of hydrogen peroxide (H_2_O_2_) 2 days after the activation of PTI, ETI^AvrRps4^ and “PTI + ETI^AvrRps4^”, respectively. The non-virulent *Pst* DC3000 *hrcC* mutant (*hrcC*^*-*^) was used to induce PTI. Using diaminobenzidine (DAB) staining, we found “PTI + ETI^AvrRps4^”, but not PTI or ETI^AvrRps4^ alone, results in H_2_O_2_ accumulation at this timepoint (Fig 1C).

### Additional physiological hallmarks of PTI are enhanced by ETI

Callose deposition is another well-documented PTI response. H_2_O_2_ promotes peroxidase-mediated cross-linking of proteins and phenolics in callose cell wall appositions during PTI^14,15^. We found that ETI^AvrRps4^ alone can also induce callose deposition. More strikingly, the overall quantity of callose deposition induced by co-activation of PTI and ETI^AvrRps4^ is statistically significantly stronger than the sum of those induced by PTI and ETI^AvrRps4^ alone (Fig 1D and E). This indicates that upon coactivation, PTI and ETI can function synergistically and mutually potentiate callose deposition.

We examined the expression of PTI-responsive genes during PTI, ETI and “PTI + ETI^AvrRps4^”. The expression of *FRK1, NHL10, FOX1* and other PTI-responsive genes is significantly higher at 24 h after “PTI + ETI^AvrRps4^” treatment compared to PTI or ETI^AvrRps4^ treatment alone (Fig 1F-H and Extended Data Fig 2P-R). In summary, PTI-induced physiological changes, such as ROS production, callose deposition and PTI-responsive gene expression, are all potentiated and enhanced by ETI.

### Activation of PTI signaling components is enhanced by ETI

We investigated how ETI potentiates PTI-induced cellular changes including ROS production. Upon PAMP recognition by PRRs, phosphorylation of the RLCK-VII family member BIK1 leads to phosphorylation of the NADPH oxidase RbohD at its 39^th^ and 343^rd^ serine residues (S39 and S343), resulting in NADPH oxidase activation. Activated RbohD catalyzes extracellular ROS production^16,17^. PTI also activates MAPKs, such as MPK3 and MPK6, which in part leads to transcriptional reprogramming^18^ (Fig 2A).

**Fig. 2.**
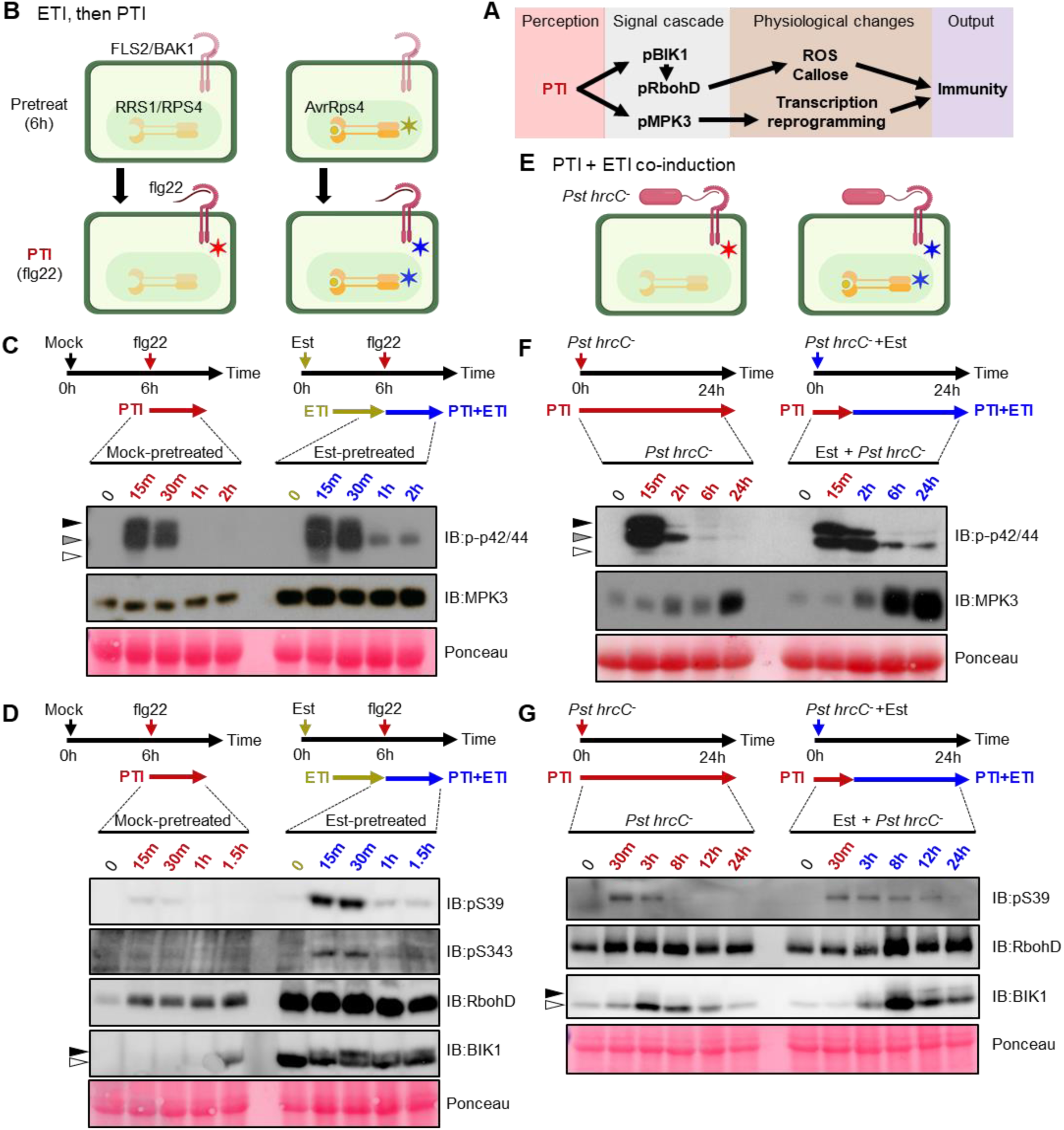
PTI-induced signaling cascade is potentiated by ETI^AvrRps4^. (A) PTI signaling pathway. (B) Schematic representation of “ETI, then PTI” experimental design for (C) and (D), ✶ indicates activated immune system. (C) ETI^AvrRps4^ pre-treatment leads to accumulation and prolonged phosphorylation of MPK3 compared to mock pre-treatment. (D) ETI^AvrRps4^ pre-treatment leads to accumulation and prolonged phosphorylation of BIK1 and RbohD (S39 and S343) compared to mock pre-treatment. (E) Schematic representation of “PTI + ETI co-induction” experimental design for (F) and (G). (F) “PTI + ETI^AvrRps4^” leads to a stronger MPK3 accumulation and prolonged phosphorylation compared to PTI. (G) “PTI + ETI^AvrRps4^” leads to a stronger BIK1 and RbohD accumulation and prolonged phosphorylation compared to PTI. For (D) and (G), microsomal fraction from the samples were isolated for immunoblotting. Arrows in (C) and (F) indicate the corresponding MAP kinases (black: pMPK6, grey: pMPK3, white: pMPK4/11). Arrows in (D) and (G) indicate the phosphorylation of BIK1 (black: pBIK1, white: BIK1). Ponceau staining was used as loading control.

We compared the induced activation of BIK1, RbohD and MAPKs during PTI to activation during “PTI + ETI^AvrRps4^”. Pre-activation of ETI^AvrRps4^ results in a prolonged flg22-induced phosphorylation of BIK1, RbohD (at S39 and S343) and MPK3 (Fig 2B-D). Similarly, co-activation of “PTI + ETI^AvrRps4^” leads to prolonged activation of BIK1, RbohD and MPK3 in response to *hrcC*^*-*^ (Fig 2E-G). In contrast, ETI^AvrRps4^ activation alone does not lead to the phosphorylation of RbohD and MAPKs (Extended Data Fig 1D and E)^19^, hence the prolonged activation of PTI signaling components is not due to the additive effect of ETI on PTI, but the potentiation of PTI by ETI. Since PTI components are turned over during activation^20–22^, replenishment of PTI components upon ETI activation would be expected to prolong and strengthen a PTI response.

### ETI leads to upregulation of PTI signaling components

To investigate how ETI potentiates PTI, we monitored accumulation of both total and activated forms of BIK1, RbohD and MPK3 proteins during “PTI + ETI^AvrRps4^” compared to PTI alone (Fig 2C, D and F-G). We also monitored protein levels of PTI signaling components during ETI activated by four additional inducible effector-expressing lines: AvrRpp4, AvrRpt2, AvrRpm1 and AvrPphB^23–26^, which are recognized by TIR-NLR RPP4 and CC-NLRs RPS2, RPM1 and RPS5, respectively^27^. In all cases, ETI elevates the protein accumulation and activation of BAK1, BIK1, RbohD and MPK3 (Fig 3A) but not of MPK6 and FLS2 (Fig 3A and Extended Data Fig 3A), consistent with the observation that PTI-induced phosphorylation of MPK3 but not MPK6, is enhanced by ETI^AvrRps4^ (Fig 2C and F).

**Fig. 3.**
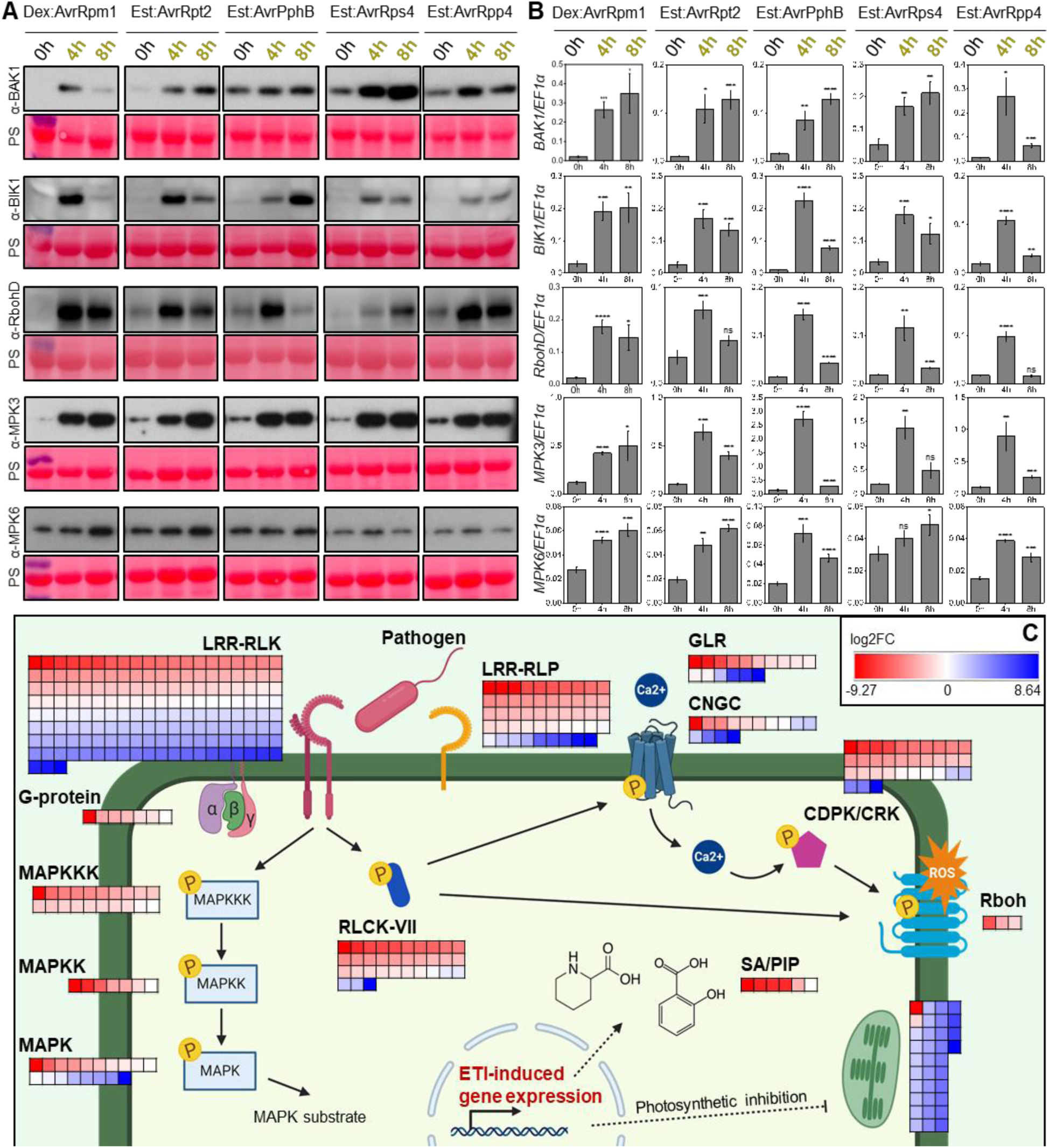
Accumulation of PTI signaling components during ETI. (A) Protein level of BAK1, BIK1, RbohD and MPK3, but not MPK6, accumulates at 4 to 8 h after ETI induction. This applies for RPM1-, RPS2-, RPS5-, RRS1/RPS4- and RPP4-induced ETI. Ponceau staining were used as loading control. (B) Relative transcript level of *BAK1, BIK1, RbohD* and *MPK3* and *MPK6*, also accumulates at 4 to 8 h after ETI induction. Student’s t-test was used to analyze significance in differences of 4 h, 8 h data points from 0 h. (ns, not significant; *; P ≤ 0.05; **, P ≤ 0.01; ***, P ≤ 0.005; ****, P ≤ 0.001). (C) RNAseq confirms the upregulation of PTI signaling pathway during ETI^AvrRps4^. Heatmap representing the expression level of PTI signaling pathway genes, defense-related hormone salicylic acid (SA) and secondary metabolite pipecolic acid (PIP) biosynthesis pathway genes and photosynthetic pathway genes at 4 h after ETI^AvrRps4^ induction. Red represents upregulation and blue represents downregulation.

Transcription and translation are strongly correlated during ETI^28,29^. We tested if elevated protein levels of PTI signaling components results from their transcriptional induction. ETI induction elevates mRNA transcript abundance of *BAK1, BIK1, RbohD, MPK3* and *MPK6* (Fig 3B). We also detected transcriptional elevation of *FLS2, MPK4, RbohF* and *ICS1* (Extended Data Fig 3B-E) indicating that ETI alone can boost the transcription of many other genes involved in PTI signaling and defense-related pathways. We therefore performed genome-wide expression profiling 4 h after the induction of ETI^AvrRps4^ and found ∼10% of the transcriptome shows significant differential gene expression upon activation of ETI^AvrRps4^ (Fig 3C, Extended Data Fig 4A and Supplementary Data 1). The majority of upregulated genes are enriched in biological processes implicated in immune responses, especially PRR signaling pathways (Fig 3C, Extended Data Fig 4A and Table 1). In addition to the tested genes, many other PTI signaling components, such as *EFR, PEPR1/2, LYK5, CERK1, XLG2, CNGC19* and *MKK4/5* are highly upregulated during ETI^AvrRps4^ (Extended Data Table 1). Genes encoding enzymes that are required for the biosynthesis of defense-related phytohormone and secondary metabolites, are also upregulated by ETI^AvrRps4^ (Extended Data Table 1)^30–37^. Overall, we conclude that ETI increases defense strength via transcriptional induction that elevates the abundance of PTI signaling components.

### ETI functions through PTI

Whether ETI and PTI activate the same or distinct mechanisms remains poorly defined, because the processes activated by ETI in the absence of PTI were rarely investigated. The data above led us to test the hypotheses that (i) PTI provides the main defense mechanism against pathogens and (ii) ETI functions primarily to enhance PTI by replenishing PTI components and restoring effector-attenuated PTI.

We challenged plants with non-virulent *hrcC*^*-*^ and found that protein levels of BIK1 and RbohD are slightly elevated during PTI, and MAPKs are activated, as indicated by their elevated phosphorylation (Extended Data Fig 5A). After infiltration with virulent strain *Pst* DC3000, PTI-induced protein accumulation of BIK1 and RbohD, and MAPK activation, are reduced compared to *hrcC*^*-*^ (Extended Data Fig 5A), consistent with disease susceptibility caused by DC3000, a process designated effector-triggered susceptibility (ETS)^1^. We co-infiltrated plants with DC3000 and estradiol to co-induce ETI^AvrRps4^ and found restoration of elevated protein levels of BIK1, RbohD and MPK3 and prolonged activation of MAPKs (Extended Data Fig 5A). This indicates that ETI activation restores PTI capacity and can overcome ETS.

During natural infections, ETI is rarely activated without PTI. We therefore propose, in a refinement of previous models^1^, that ETI provides robust resistance by restoring and elevating abundance of PTI signaling components, compensating for their turnover upon activation and attenuation by ETS (Extended Data Fig 5B). This model also implies that NLR-mediated resistance functions through PTI. We therefore tested the requirement of PTI in NLR-dependent resistance by infiltrating the PTI-compromised co-receptor double mutant *bak1-5 bkk1-1* with DC3000 delivering AvrRps4. Remarkably, this mutant is as susceptible as the *rps4-21 rps4b-1* mutant that cannot detect AvrRps4 (Figure 3A), demonstrating that PTI is required for *RPS4/RRS1*-dependent resistance to bacteria. This shows that activation of ETI alone in the absence of PTI is not sufficient to provide effective resistance against *P. syringae* in Arabidopsis. In addition, Yuan *et al* (BIORXIV/2020/031294) provide complementary data, independently showing that PTI is required for bacterial resistance mediated by multiple NLRs.

### PTI potentiates ETI-induced hypersensitive response

ETI in the presence of PTI often culminates in a hypersensitive cell death response (HR). When Arabidopsis is infiltrated with a non-pathogenic strain *P. fluorescens* Pf0-1 delivering wild-type AvrRps4 (Pf0-1:AvrRps4^WT^), which triggers “PTI + ETI^AvrRps4^”, cell death is observed (Extended Data Fig 6A). However, ETI^AvrRps4^ alone does not lead to HR (Extended Data Fig 6A)^12^. To explain this discrepancy, we hypothesized that co-activation of PTI is required for ETI^AvrRps4^ to induce HR.

We used a Pf0-1 strain delivering a non-functional mutant allele of AvrRps4 (Pf0-1:AvrRps4^mut^) to activate PTI^38^, and estradiol in the inducible AvrRps4 line to activate ETI^AvrRps4 12^. Co-infiltration of Pf0-1:AvrRps4^mut^ and estradiol in the inducible AvrRps4 line leads to the co-activation of PTI and ETI^AvrRps4^ (“PTI + ETI^AvrRps4^”), which results in similar macroscopic HR induced by Pf0-1:AvrRps4^WT^ and a stronger electrolyte leakage (a widely used indicator of cell death) compared to PTI or ETI^AvrRps4^ alone (Extended Data Fig 6A and B). To test if other PTI-inducers could also potentiate cell death, we repeated the estradiol co-filtration experiment with either *hrcC*^*-*^, a wild-type Pf0-1 strain without type-III secretion system or a mixture of PAMPs flg22, elf18 and a danger-associated molecular pattern (DAMP) ligand pep1 to activate PTI^2^. In all cases, PAMP co-infiltration combined with estradiol induction of AvrRps4 resulted in HR (Figure 4B). This implies that PTI potentiates ETI-activated HR.

**Fig. 4.**
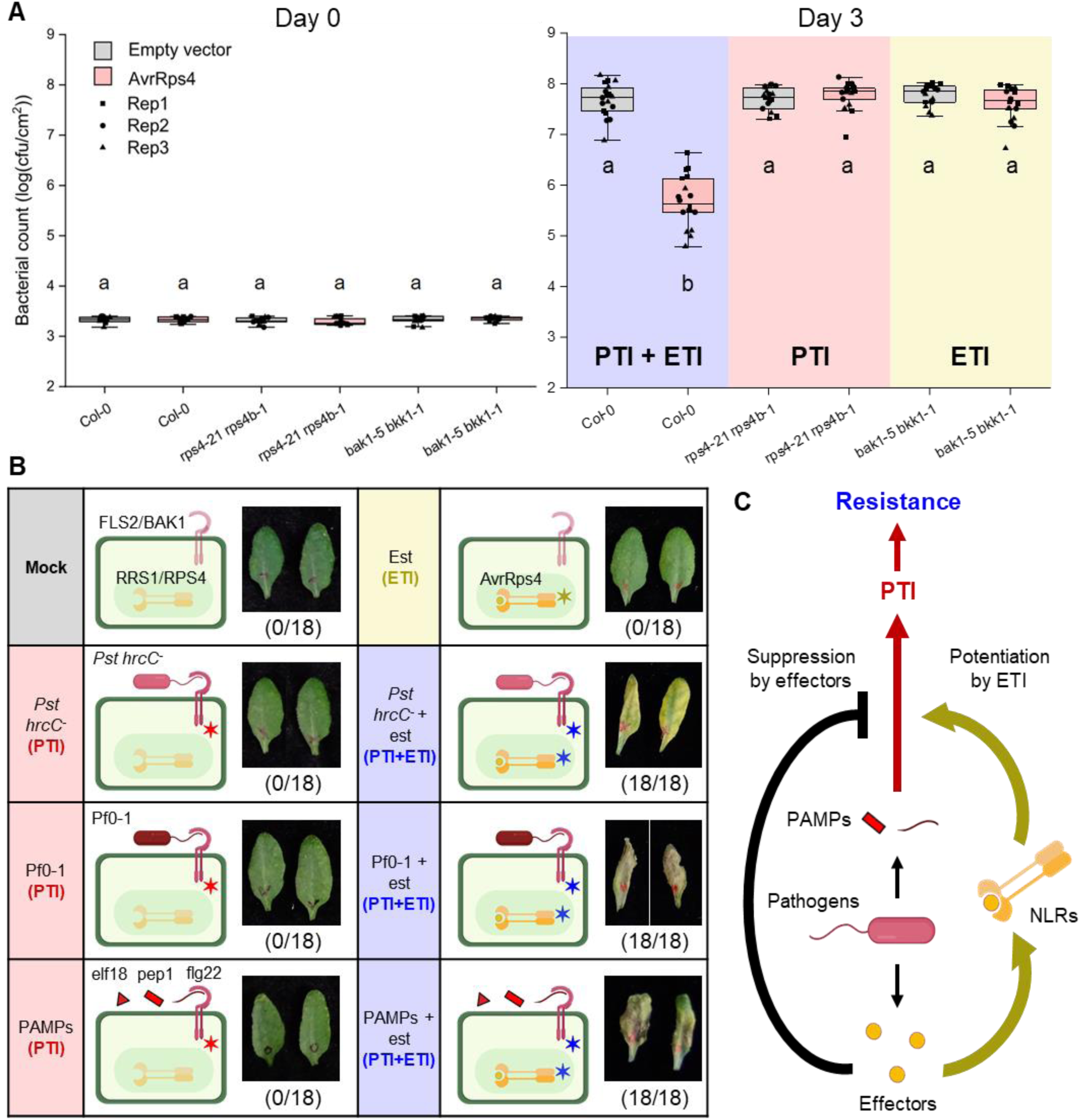
PTI and ETI function synergistically to provide robust immunity. (A) Both PTI and ETI are required to provide effective immunity against *Pseudomonas syringae*. Col-0, *rps4-21 rps4b-1* and *bak1-1 bkk1-1* were infected with *Pseudomonas syringae* pv. *tomato* strain DC3000 (*Pst*) carrying empty vector (control) or AvrRps4. Both *rps4-21 rps4b-1* (PTI only) and *bak1-1 bkk1-1* (ETI only) are insufficient to provide resistance against *Pst*:AvrRps4 compared to Col-0 (PTI + ETI). (B) ETI^AvrRps4^ alone does not lead to macroscopic HR. Together with PTI, activated by either avirulent *Pst hrcC*^*-*^, *Pseudomonas fluorescens* (Pf0-1) or mixture of flg22, elf18 and pep1 (PAMPs), ETI leads to macroscopic HR. ✶ in the schematic diagrams indicate activated immune system. (C) Schematic representation of the plant immune system. PAMPs from pathogens are recognized by plant PRRs and induce PTI (red). Virulent pathogen secretes effectors to suppress PTI (black). Effectors are recognized by NLRs and induce ETI (dark yellow), which restores and potentiates PTI to produce robust immune response (blue).

Like ETI^AvrRps4^, ETI^AvrRpp4^ alone cannot induce HR cell death, but co-activation of PTI and ETI^AvrRpp4^ can (Extended Data Fig 7A, B). Both effectors are recognized by TIR-NLRs. In contrast, inducible expression of AvrRpt2, AvrRpm1 and AvrPphB that are recognized by CC-NLRs can trigger cell death in the absence of PTI in Arabidopsis (Extended Data Fig 7A). Recently, it was reported that upon activation, the CC-NLR ZAR1 forms a pentameric “resistosome” that induces cell death dependent on oligomerization of its N-terminal domains (NTDs)^7,39^, whereas activation of defense by TIR-NLRs requires the NADase activities of their NTDs^9^. Thus, the differential HR cell death induced by TIR-NLRs and CC-NLRs reported here might be due to their different NTD-specific signaling mechanisms. By reducing levels of estradiol or dexamethasone, we defined sub-lethal levels of AvrRpt2, AvrRpm1 and AvrPphB induction. At these levels, CC-NLR mediated HR cell death was also enhanced by PTI co-activation (Extended Data Fig 7B). Thus, with all NLRs in Arabidopsis that we examined, we found that PTI activation enhances ETI-induced HR cell death.

### PTI potentiates ETI-induced HR through MAPKs and NADPH oxidases

Previously, PTI signaling components MAPKs and Rboh proteins were reported to be required by HR induced by ETI in the presence of PTI^40,41^. To investigate the molecular basis of PTI-enhanced ETI-associated HR, we investigated the role of the MAPKs and Rbohs during ETI alone. We found that MAPKs are phosphorylated during ETI^AvrRpm1^, ETI^AvrRpt2^ and ETI^AvrPphB^, but not during ETI^AvrRps4^ or ETI^AvrRpp4^ (Extended Data Fig 1D and 8A-D). However, none of the inducible ETIs tested led to RbohD phosphorylation at S39 (Extended Data Fig 1E and 8E-I), consistent with our observation that ETI alone does not activate a strong ROS burst (Fig 1A-B). ETI^AvrRpt2^ leads to RbohD phosphorylation at S343 and S347^42^, which might explain why ETI^AvrRpt2^ activates a weak ROS burst (Extended Data Fig 2H-I).

Since ETI potentiates the PTI-induced activation of MPK3, and ETI alone leads to weak or no activation of these signaling components, we tested if the enhancement of HR by PTI involves the ETI-potentiated activity of MPKs. To test this, we used an Arabidopsis line *MPK6SR*, in which an *mpk3 mpk6* double mutant is complemented by a mutant MPK6 allele (MPK6^YG^) whose activity can be inhibited by a bulky protein kinase inhibitor, the ATP analogue 1-NA-PP1,which does not inhibit wild-type MAP kinases^43^. We tested the response to Pf0-1:AvrRps4^WT^ (“PTI + ETI^AvrRps4^”) in the *MPK6SR* line in the presence or absence of 1-NA-PP1. Consistent with previous reports^40^, we found that inhibition of MPK6^YG^ in *MPK6SR* prevents ETI^AvrRps4^-associated HR even in the presence of PTI (Extended Data Fig 9), showing that MAP kinase activation makes an indispensable contribution to the HR.

## Discussion

Although PTI mechanisms have been defined in detail, the relationship between PTI and ETI, and the consequences of activating ETI without PTI, were poorly understood. Most studies have compared PTI with “PTI + ETI”, and there are few studies on ETI alone in the absence of PTI. Our data show that PTI and ETI result in entirely distinct physiological outputs and initiate distinct chains of events. We also discovered that the stronger immune response during “PTI + ETI” is due to mutual potentiation of these two systems.

Importantly, by showing that ETI requires PTI to provide effective resistance, we shed new light on how ETI halts pathogens. PTI has been shown to halt pathogens through nutrient restriction, cell wall fortification, suppression of bacterial type III secretion and induction of antimicrobial compounds^14,15,44–47^. Our work implies that ETI halts pathogens through the potentiation of PTI. The mechanism(s) by which ETI leads to increased transcript and protein abundance of PTI signaling components remain to be determined.

We provide new insights into how surface receptor-mediated immunity and intracellular receptor-mediated immunity work together to provide a more robust disease resistance than either alone. Our data, and those of Yuan *et al* (BIORXIV/2020/031294), support a model in which cellular processes activated by cell-surface PRR-dependent signaling are the primary source of immunity, and intracellular NLR receptors act to replenish PRR signaling components and enhance PRR-dependent signaling, counteracting its attenuation by turnover upon activation and by effector-dependent mechanisms that suppress host immunity (Fig 4C). In turn, cell-surface receptor-mediated immunity can potentiate immune responses activated by intracellular receptors, such as HR cell death to further restrict the spread of pathogens. These data are highly relevant to elevating crop disease resistance. Many NLR-encoding genes are semi-dominant, suggesting ETI strength is rate-limiting for resistance. Thus, stacks of multiple NLR-encoding genes should provide physiologically stronger resistance, as well enhancing genetic durability, and are a potential source of non-host resistance^48^. Other reports have indicated synergistic functions of cell-surface and intracellular receptors in mammalian immunity^49–53^, highlighting the relevance of these insights to multiple host-pathogen systems.

## Methods

### Plant material and growth conditions

*Arabidopsis thaliana* accessions and Columbia-0 (Col-0) were used as wild type in this study. Seeds were sown on compost and plants were grown at 21°C with 10 h under light and 14 h in dark, and at 70% humidity. The light level is approximately 180-200 µmols with fluorescent tubes. Information about all plant materials can be found in the referred literatures^12, 23-26, 40, 54-56^, and some of which were kindly provided by Jeffery Dangl (Department of Biology, The University of North Carolina at Chapel Hill), Roger Innes (Department of Biology, Indiana University), Shuta Asai (RIKEN, Japan), Shuqun Zhang (Division of Biochemistry, University of Missouri), Xiufang Xin (Shanghai Institutes for Biology Sciences, Chinese Academy of Sciences) and Cyril Zipfel (The Sainsbury Laboratory, UK).

### Immunoblotting

*Arabidopsis thaliana* leaves were infiltrated with different treatment solution as described. Samples were collected at indicated time points and snap-frozen in liquid nitrogen. Arabidopsis seedlings were grown for 10 days after germination and were treated with different treatment solution with indicated time points and snap-frozen in liquid nitrogen. Samples were lysed and proteins were extracted using GTEN buffer (10% glycerol, 25 mM Tris pH 7.5, 1 mM EDTA, 150 mM NaCl) with 10 mM DTT, 1% NP-40 and protease inhibitor cocktail (cOmplete™, EDTA-free; Merck), phosphatase inhibitor cocktail 2 (Sigma-Aldrich; P5726) and phosphatase inhibitor cocktail 3 (Sigma-Aldrich; P0044). After centrifugation at 13,000 rpm for 10 minutes to remove cell debris, protein concentration of each sample was measured using the Bradford assay (Protein Assay Dye Reagent Concentrate; Bio-Rad). After normalization, extracts were incubated with 2× TruPAGE™ LDS Sample Buffer (Sigma-Aldrich) at 70°C for 10 minutes. SDS-PAGE gels of different percentages were used to run protein samples of difference sizes. After transferring proteins from gels to PVDF membranes (Merck-Millipore) using Trans-Blot Turbo System (Bio-Rad), membranes were blocked with 5% nonfat dried milk in TBST for 1 h, immunoblotted with either BAK1 antibody (Agrisera; AS12 1858), BIK1 antibody (Agrisera; AS16 4030), RbohD antibody (Agrisera; AS15 2962), MPK3 antibody (Sigma-Aldrich; M8318), MPK6 antibody (Sigma-Aldrich; A7104) or Phospho-p44/42 MAPK (erk1/2) (Thr202/Tyr204) (D13.14.4E) XP® Rabbit monoclonal antibody (Cell Signaling Technology; 4370). Anti-Rabbit IgG (whole molecule)–Peroxidase antibody produced in goat (A0545; Merck-Sigma-Aldrich) was used as secondary antibody following the use of above antibody. Ponceau S solution (P7170; Sigma-Aldrich) was used to stain the PVDF membrane for loading control.

### Plasma membrane extraction for the detection of phosphorylated RbohD and BIK1

Minute™ Plant Plasma Membrane Protein Isolation Kit (Invent Biotechnologies, SM-005-P) was used to extract total membrane fraction from Arabidopsis samples. Protein concentration of the cytosolic fraction from each sample was measured using the Bradford assay (Protein Assay Dye Reagent Concentrate; Bio-Rad). After normalization, total membrane fractions were dissolved in 2× TruPAGE™ LDS Sample Buffer (Sigma-Aldrich) at 70 °C for 5 minutes (in a minimal of 80μl). 6% SDS-PAGE gels were used to run the protein samples. After transferring proteins from gels to PVDF membranes (Merck-Millipore) using Trans-Blot Turbo System (Bio-Rad), membranes were blocked with 5% nonfat dried milk in TBST for 1 h, immunoblotted with P39-RbohD or P343-RbohD antibodies kindly provided by Jian-Min Zhou (Institute of Genetics and Developmental Biology, Chinese Academy of Sciences)^17^. Anti-Rabbit IgG (whole molecule)–Peroxidase antibody produced in goat (A0545; Merck-Sigma-Aldrich) was used as secondary antibody. Ponceau S solution (P7170; Sigma-Aldrich) was used to stain the PVDF membrane for loading control.

### Gene expression measurement by reverse transcription-quantitative polymerase chain reaction (RT-qPCR)

For gene expression analysis, RNA was isolated from 5-week-old Arabidopsis leaves with RNeasy Plant Mini Kit (74904; Qiagen) and used for subsequent RT-qPCR analysis. RNA was extracted with Quick-RNA Plant Kit (R2024; Zymo Research) and treated with RNase-free DNase (4716728001; Merck-Roche). Reverse transcription was carried out using SuperScript IV Reverse Transcriptase (18090050; ThermoFisher Scientific). qPCR was performed using a CFX96 Touch™ Real-Time PCR Detection System. Primers for qPCR analysis of, *BRI1-Associated receptor Kinase 1* (*BAK1*), *Botrytis–Induced Kinase 1* (*BIK1*), *Respiratory burst oxidase homolog protein D* (*RbohD*), *Mitogen-activated Protein Kinase 3* (*MPK3*), *Isochorismate Synthase1* (*ICS1*), *Flagellin-sensitive 2* (*FLS2*), *Mitogen-activated Protein Kinase 4* (*MPK4*), *Respiratory burst oxidase homolog protein F* (*RbohF*), *Mitogen-activated Protein Kinase 6* (*MPK6*), *Flg22-induced Receptor-like Kinase 1* (*FRK1*), *NDR1/HIN1-Like protein 10* (*NHL10*), *Fad-linked OXidoreductase 1* (*FOX1*), *PERoxidase 4* (*PER4*), *probable WRKY transcription factor 31* (*WRKY31*) and *Elongation Factor 1 Alpha* (*EF1α*) are listed in Supplementary Information. Data were analyzed using the double delta Ct method^57^. A summary of statistical analysis can be found in Supplementary Data 2.

### ROS burst assay

Leaf discs harvested with a 6-mm-diameter cork borer from 5-week-old plants were placed in 96-well plates with 200 µL of deionized water overnight in dark. 200 µL of 20 mm luminol (Sigma-Aldrich, A8511), 0.02 mg/mL horseradish peroxidase (Sigma-Aldrich, P6782) and indicated elicitors were added in each well. ROS production was measured with a Photek camera (East Sussex, UK). Data from each treatment is represented by 40 leaf discs in one biological replicate. Every plate was measured over the indicated time. A summary of statistical analysis can be found in Supplementary Data 2.

### DAB staining

3,3’-diaminobenzidine (Sigma-Aldrich, D8001) was dissolved in pH 3.8 water (1 mg/mL) and the pH is adjusted to 6. Arabidopsis leaves following indicated treatment were vacuum infiltrated in DAB solution for 30 minutes and incubated in room temperature for 2 h. The DAB solution was replaced with 100% ethanol and then boiled for 1 minute. The leaves are then further de-stained with 70% ethanol. De-stained leaves were scanned with EPSON Perfection V600 Photo. Scale bar = 0.5 cm.

### Callose quantification

Leaves from 5-week-old Arabidopsis were hand-infiltrated with the indicated solution and covered for 24 h. Leaves were then hand-infiltrated with 1× PBS buffer containing 0.01% Aniline Blue. Leaf discs were then harvested with a 6-mm-diameter cork borer for imaging. Images were taken by an epifluorescence microscope with UV filter (excitation, 365/10 nm; emission, 460/50 nm). The number of callose dots was calculated by ImageJ software. One leaf disc was harvest per leaf.

At least 7 leaves from individual plants were included per treatment in one biological replicate. A summary of statistical analysis can be found in Supplementary Data 2.

### HR assay in Arabidopsis

*Pseudomonas fluorescens* engineered with a type III secretion system (Pf0-1 ‘EtHAn’ strains) expressing effectors, AvrRps4, AvrRps4^KRVY135-138AAAA^ (mutant AvrRps4), AvrRpm1, AvrRpt2, AvrPphB or pVSP61 empty vector were grown on selective KB plates for 24 h at 28°C. Wild-type *Pseudomonas fluorescens* were grown on KB plates with chloramphenicol for 24 h at 28°C. *Pseudomonas syringae pv. tomato* strain DC3000 *hrcC*^*-*^ or DC3000 were grown on KB plates with kanamycin for 48 h at 28°C. Bacteria were harvested from the plates, resuspended in infiltration buffer (10 mM MgCl_2_) and the concentration was adjusted to indicated OD600 (supplementary table 2). The abaxial surfaces of 5-week-old Arabidopsis leaves were hand infiltrated with indicted solution by a 1-ml needleless syringe. Cell death was monitored at indicated time points after infiltration.

### Electrolyte leakage assay

Indicated solutions were hand infiltrated in 5-week-old Arabidopsis leaves with a 1-ml needleless syringe for electrolyte leakage assay. Leaf discs were taken with a 2.4-mm-diameter cork borer from infiltrated leaves. Discs were dried and washed in deionized water for 1 h before being floated on deionized water (15 discs per sample, three samples per biological replicate). Electrolyte leakage was measured as water conductivity with a Pocket Water Quality Meters (LAQUAtwin-EC-33; Horiba) at the indicated time points. A summary of statistical analysis can be found in Supplementary Data 2.

### RNAseq and data analysis

Leaves from 5-week-old Arabidopsis est:AvrRps4 or est:AvrRps4^mut 19^ were hand-infiltrated with 50 μM estradiol for 0 or 4 h. Samples were collected and total RNA was isolated with TRI Reagent^®^ (T9424: Sigma-Aldrich) and RNA Clean & Concentrator-25 Kit (R1018; Zymo Research). RNA samples are processed by BGI and libraries are sequenced with BGISEQ-500 sequencing platform. At least 10 M single-end 50-bp reads are obtained for each RNAseq library. Adaptor-trimmed clean reads have been uploaded to the European Nucleotide Archive (ENA) (publica accessible number is available upon acceptation for publication). After FastQC, we used Kallisto to map and quantify our RNAseq reads^58^, and kallisto_quant output files are submitted to the 3-D RNAseq tool for statistics and data visualization^59^.

### Statistical data analysis

Statistical data were analyzed using the R software, and the data were plotted using the Origin software. Statistical analysis was performed with one-way ANOVA analysis followed by Tukey’s post-hoc HSD test for all experiments, except for qPCR assays. Data points with different letters indicate significant differences of P<0.01 for Tukey’s HSD test results. Data points are plotted onto the graph, and number of samples for each data are indicated in corresponding figure legends. Three biological replicates were tested, and individual biological replicates are indicated with different shapes of the data points. qPCR assay results were analyzed using Student’s T-test for statistical significance (ns, not significant; *; P ≤ 0.05; **, P ≤ 0.01; ***, P ≤ 0.005; ****, P ≤ 0.001) between samples. A summary of statistical analysis can be found in Supplementary Data 2.

## Acknowledgments

We thank Jeff Dangl, Shuta Asai, Roger Innes, Kenichi Tsuda, Shuqun Zhang, Cyril Zipfel, Jian-Min Zhou, Brian Staskawicz, Cecilia Cheval, Yasuhiro Kadota, Ruby O’Grady, Maddy Morris, Jack Rhodes and Yuli Ding for providing materials, discussion and technical support. We also thank Cyril Zipfel, Minhang Yuan, Xiufang Xin and Sheng-Yang He for critical reading of the manuscript. We thank the Gatsby Foundation (United Kingdom) for funding to the JDGJ laboratory. BN was supported by the Norwich Research Park Biosciences Doctoral Training Partnership from the Biotechnology and Biological Sciences Research Council (BBSRC) (grant agreement: BB/M011216/1). HA was supported by European Research Council Advanced Grant “ImmunitybyPairDesign” (grant agreement: 669926). PD acknowledges support from the European Union’s Horizon 2020 Research and Innovation Program under Marie Sklodowska-Curie Actions (grant agreement: 656243) and a Future Leader Fellowship from BBSRC (grant agreement: BB/R012172/1).

## Author contributions

BN, PD and JDGJ conceptualized the study. Experiments are carried out by BN and HA. Data analysis was performed by BN, HA and PD. BN and PD wrote the manuscript. BN, HA, PD and JDGJ reviewed and edited the manuscript.

## Competing interest declaration

The authors declare no competing interests.

## Data and materials availability

All data needed to replicate the work are present in materials and supplementary information.

## Additional Information

Supplementary Information is available for this paper.

## Extended Data

**Extended Data Fig. 1.**
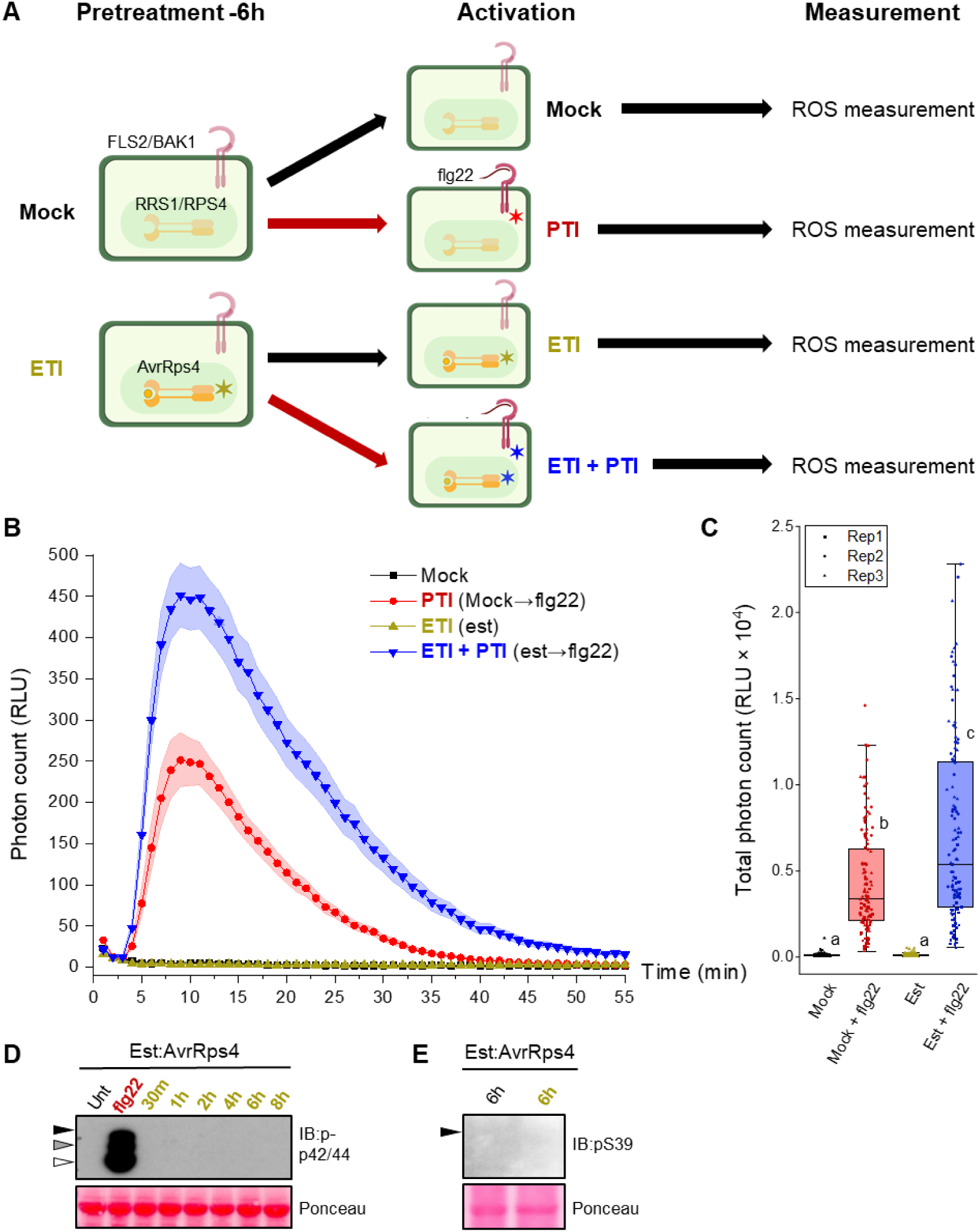
ETI^AvrRps4^ leads to enhanced ROS production triggered by PTI. (A) Schematic representation of experimental design for (B) and (C). 5-week old leaf disks from est:AvrRps4 line were soaked in mock solution (1% DMSO) or 50 μM est (estradiol, triggers ETI^AvrRps4^) for 6 h. Mock or 100 nM flg22 were then added to the leaves and ROS accumulation was measured. (B) ETI^AvrRps4^ pre-treatment leads to stronger and prolonged ROS accumulation compared to mock pre-treatment. (C) Total ROS accumulation in ETI^AvrRps4^-pretreated leaves is significantly higher than mock-pretreated leaves. Data points from 3 biological replicates were analyzed with one-way ANOVA followed by Tukey’s HSD test. Data points with different letters indicate significant differences of P < 0.01. (D-E) ETI^AvrRps4^ alone does not lead to MAPKs activation or RbohD-S39 phosphorylation (dark yellow). These two experiments were performed together with other inducible lines in extended data figure 8. For detailed experimental setup and controls refer to Extended Data Fig 8.

**Extended Data Fig. 2.**
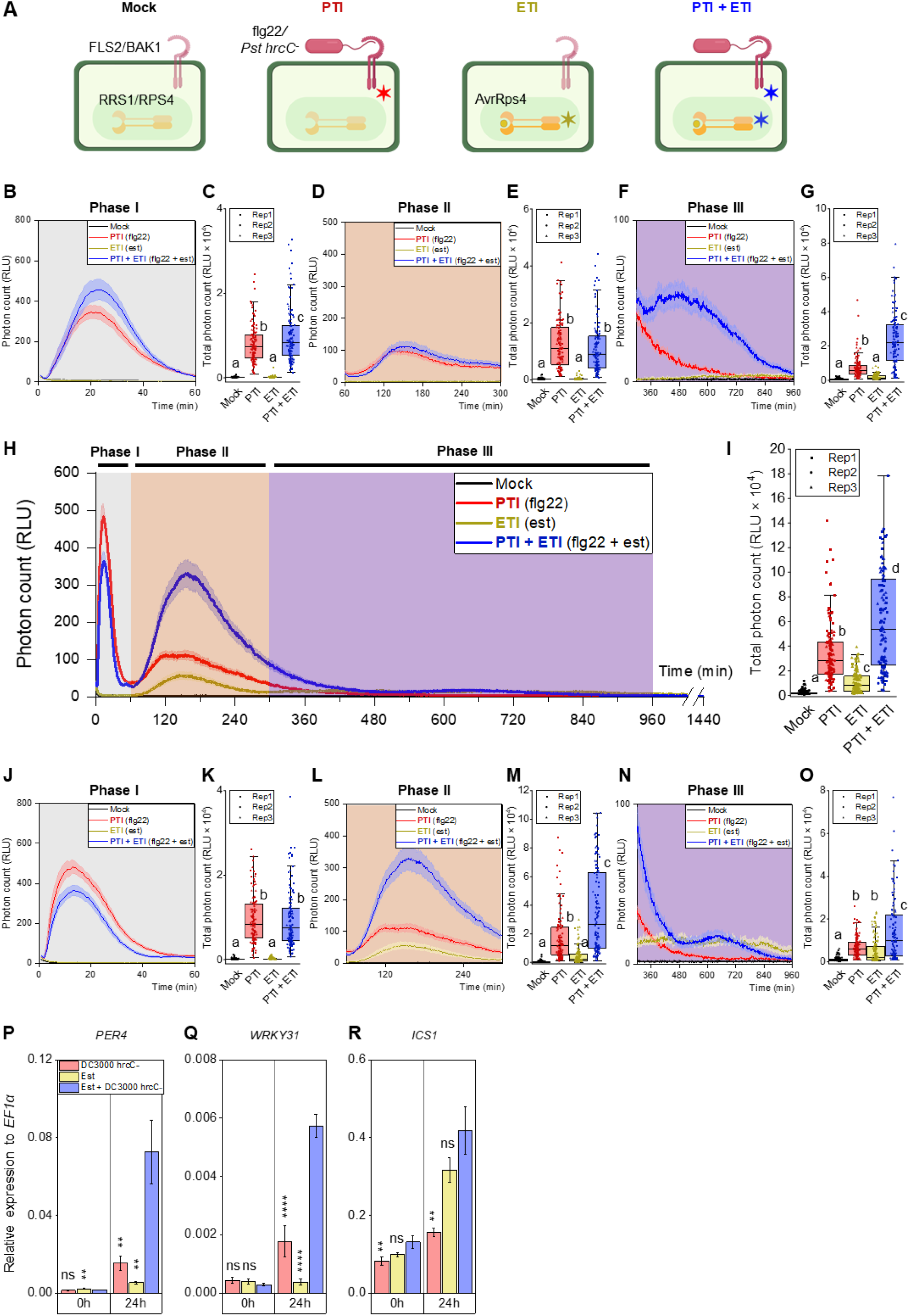
Potentiation of PTI-induced physiological changes during ETI. (A) Schematic representation of “PTI + ETI co-induction” experimental design. (B-G) “PTI + ETI^AvrRps4^” leads to enhanced ROS production from 300-960 minutes. ROS production in est:AvrRps4 during phase I (B), phase II (D) and phase III (F). Total ROS production in est:AvrRps4 during phase I (C), phase II (E), phase III (G). (H) “PTI + ETI^AvrRpt2^” leads to enhanced ROS production from 60-480 minutes. (I) Total ROS production in “PTI + ETI^AvrRpt2^” treated leaves is significantly higher than PTI-treated leaves. ROS production of est:AvrRpt2 line in phase I (J), phase II (L) and phase III (N). Total ROS production in est:AvrRpt2 line during phase I (K), phase II (M), phase III (O). Shaded curve in S2B, D, F, H, J, L and N represents S.E.. For Extended Data Fig 2C, E, G, I, K, M and O, data points from 3 biological replicates were analyzed with one-way ANOVA followed by Tukey’s HSD test. Data points with different letters indicate significant differences of P < 0.01. “PTI + ETI^AvrRps4^” leads to a stronger (P) *PER4* and (Q) *WRKY31* transcript accumulation compared to PTI or ETI^AvrRps4^ alone. (R) *ICS1* acts as a control. There is no significant difference in *ICS1* expression between PTI + ETI^AvrRps4^ and ETI^AvrRps4^ alone. The average of data points from 3 biological replicates were plotted onto the graphs, with ±S.E. for error bars. Student’s t-test was used to analyze significance differences between “PTI + ETI^AvrRps4^” and PTI or ETI^AvrRps4^. (ns, not significant; *; P ≤ 0.05; **, P ≤ 0.01; ***, P ≤ 0.005; ****, P ≤ 0.001).

**Extended Data Fig. 3.**
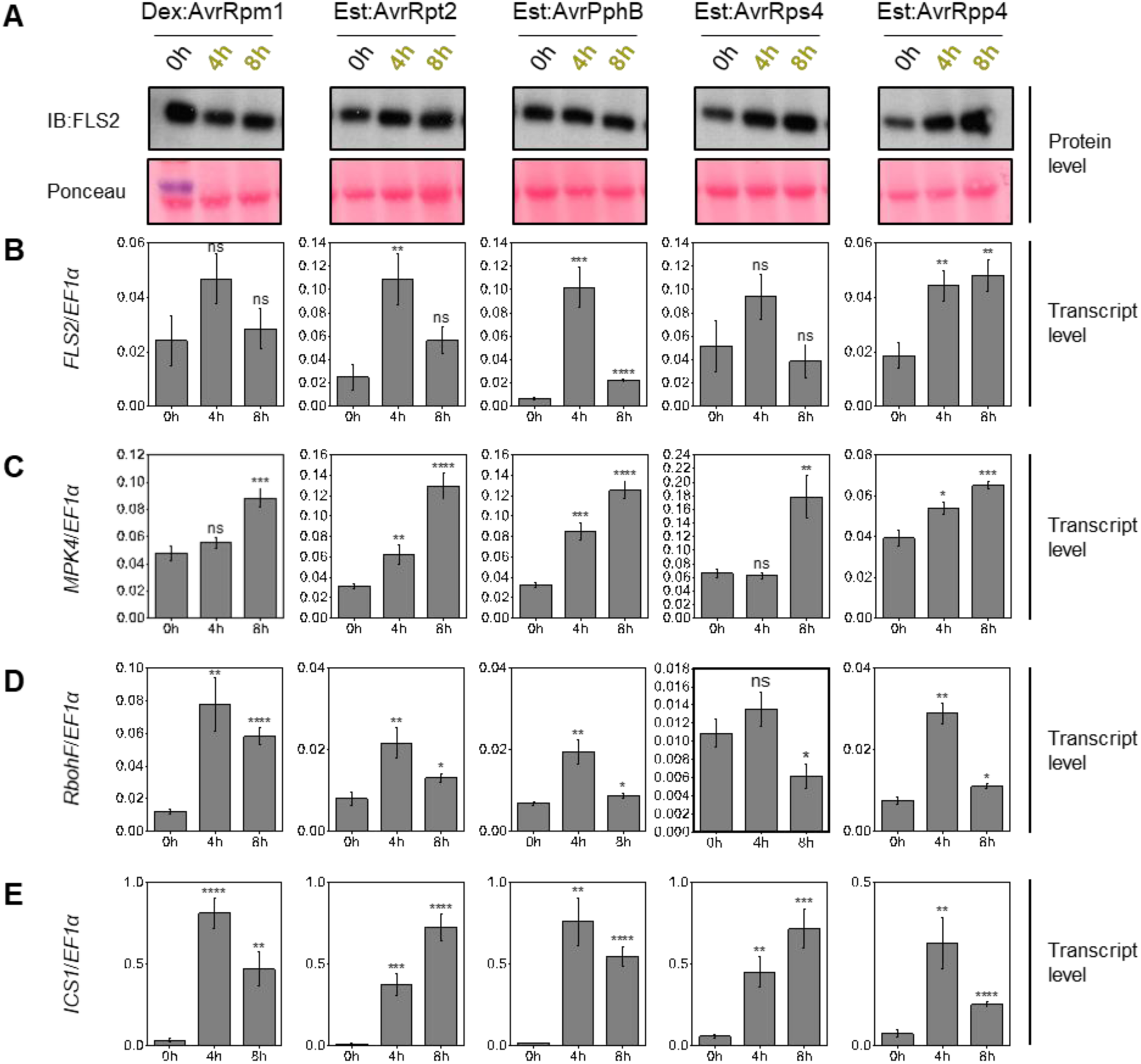
Protein and transcript levels of multiple genes during ETI. FLS2 protein (A) and transcript level (B) during ETI. Transcript level of (C) *MPK4*, (D) *RbohF*, (E) *ICS1* during ETI. 5-week old leaves of inducible-AvrRpm1, AvrRpt2, AvrPphB, AvrRps4 and AvrRpp4 lines were infiltrated with 50 μM dex (for dex:AvrRpm1) or 50 μM est. Samples were collected at 0, 4 and 8 hours post infiltration (hpi) for RNA and protein extraction. Ponceau staining were used as loading control for the above experiments. Extracted RNA were analyzed by qPCR and expression level is presented as relative to *EF1α*. The average of data points from 3 biological replicates were plotted onto the graphs, with ±S.E. for error bars. Student’s t-test was used to analyze significance in differences of 4 h, 8 h data points from 0 h. (ns, not significant; *; P ≤ 0.05; **, P ≤ 0.01; ***, P ≤ 0.005; ****, P ≤ 0.001).

**Extended Data Fig. 4.**
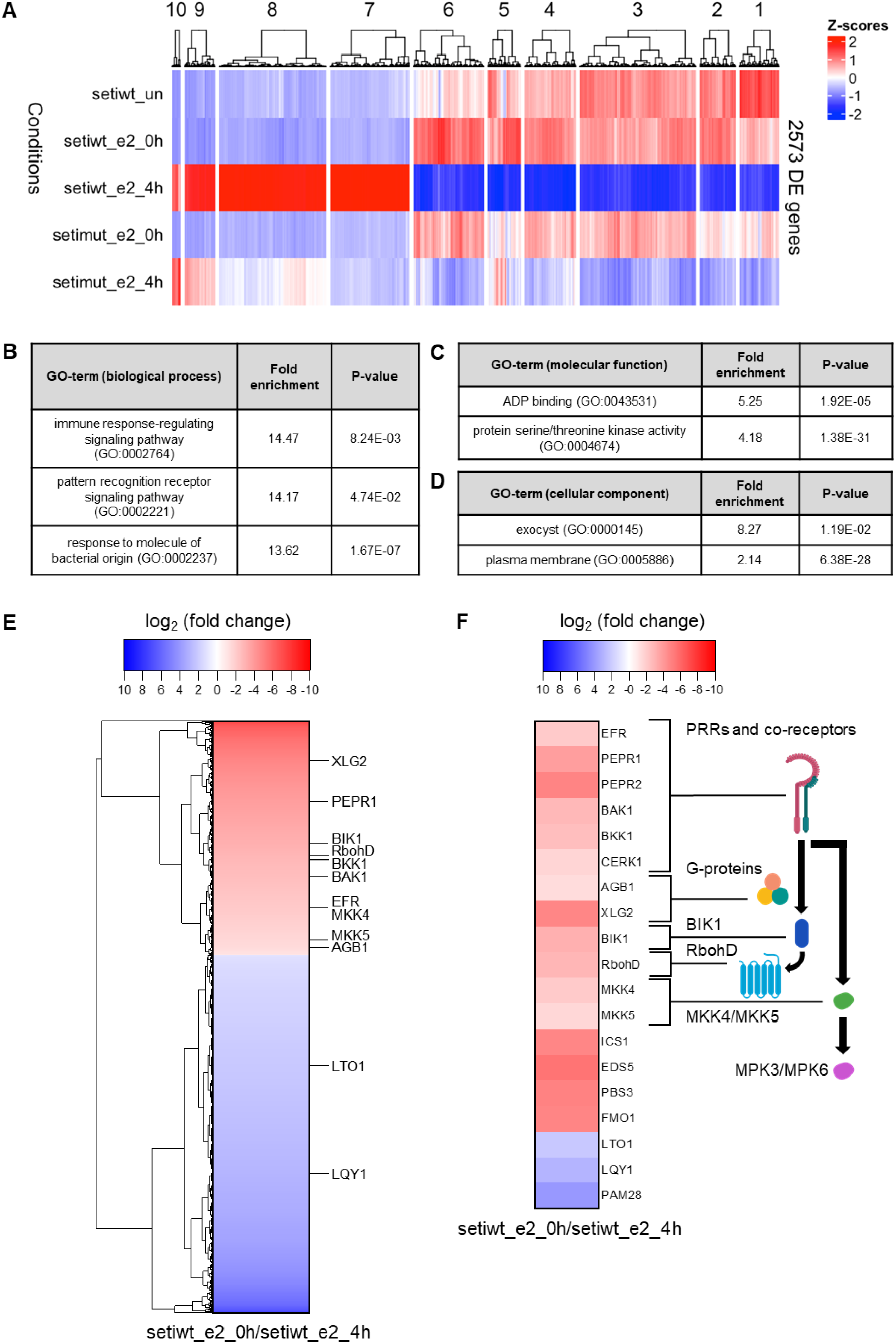
Genome-wide gene expression profiling during ETI^AvrRps4^. 2573 differential expressed (DE) genes were identified as significant in comparison between est:AvrRps4 (SETI) treated with estradiol (chemical abbreviation: E2) for 0 h (seti_e2_0h) and est:AvrRps4 treated with est for 4 h (seti_e2_4h). DE genes with adjusted p-value < 0.01 is categorized as significant. (A) Heatmap representing the 2573 DE genes during 5 treatments: est:AvrRps4 untreated (setiwt_un), est:AvrRps4 treated with est for 0 h (setiwt_e2_0h), est:AvrRps4 treated with est for 4 h (setiwt_e2_4h), est:AvrRps4^mut^ (inducible AvrRps4^KRVY135-138AAAA^ mutant) treated with est for 0 h (setimut_e2_0h) and est:AvrRps4^mut^ treated with est for 4 h (setimut_e2_4h). Genes that are specifically upregulated during ETI^AvrRps4^ are in cluster 7 and 8. (B-D) GO enrichment analysis of genes from cluster 7 and 8. (B) Top three significantly enriched biological process GO-terms in cluster 7 and 8. (C) Top two significantly enriched molecular function GO-terms in cluster 7 and 8. (D) Top two significantly enriched cellular component GO-terms in cluster 7 and 8. For details of GO enrichment analysis refer to materials and methods section. (E) Heatmap representing the expression level of 2573 DE genes during seti_e2_0h relative to seti_e2_4h. (F) Heatmap representing the expression level of PTI signaling pathway genes, SA biosynthesis pathway genes and photosynthetic pathway genes during seti_e2_0h relative to seti_e2_4h. ETI^AvrRps4^ leads to the upregulation of genes in PTI signaling pathway and the downregulation of genes in photosynthetic pathway. (E-F). Red (negative log2(fold change)) represents genes that are significantly induced during seti_e2_4h compared to seti_e2_0h. Blue (positive log2(fold change)) represents genes that are significantly repressed during seti_e2_4h compared to seti_e2_0h. For detailed statistical analysis refer to Methods section. For full list of DE genes refer to Supplementary Data 1.

**Extended Data Table 1.** Expression of level of PTI signaling pathway genes, Phytohormone biosynthesis pathway genes (SA/PIP) and photosynthetic pathway genes 4 h after ETI^AvrRps4^ induction. Red (negative log2FC (fold change)) represents genes that are significantly induced and blue (positive log2FC) represents genes that are significantly repressed. Adjusted p-value (adj.pval) < 0.05 is considered as significant. For full list of DE genes refer to Supplementary Data 1.

| Target | log2FC | adj.pval |
| --- | --- | --- |
| LRR-RLKs |  |  |
| *PEPR2* | -4.8116 | 0.002686 |
| *NILR1* | -4.51897 | 0.004565 |
| *PEPR1* | -3.81316 | 0.007323 |
| *RLK7* | -3.05215 | 0.005394 |
| *BAK1* | -2.79654 | 0.00124 |
| *BKK1* | -2.56175 | 0.004736 |
| *FLS2* | -2.31705 | 0.017503 |
| *PSKR1* | -2.28895 | 0.001906 |
| *EFR* | -2.08306 | 0.004727 |
| *SOBIR1* | -1.50852 | 0.024598 |
| *BAM1* | 1.891965 | 0.035242 |
| *IRK* | 2.535638 | 0.016176 |
| *ERECTA* | 2.767986 | 0.003164 |
| *RDK1* | 3.930124 | 0.006537 |
| *BAM2* | 4.00554 | 0.002686 |
| *ERECTA-LIKE2* | 4.966724 | 0.039776 |
| LRR-RLPs |  |  |
| *RLP30* | -3.8728 | 0.007542 |
| *CLV2* | 1.499033 | 0.047117 |
| Other PRRs |  |  |
| *LYK2* | -4.30363 | 0.042594 |
| *LORE* | -3.25007 | 0.000782 |
| *LYK5* | -2.51618 | 0.029581 |
| *CERK1* | -1.64638 | 0.004578 |
| RLCK-VII |  |  |
| *BIK1* | -3.10121 | 0.00196 |
| *PBL40* | -1.84141 | 0.002449 |
| *PBL39* | -1.64469 | 0.001972 |
| G-protein |  |  |
| *XLG2* | -4.7505 | 0.001035 |
| *GPA1* | -1.62413 | 0.005623 |
| *AGB1* | -1.36574 | 0.00495 |
| CPKs |  |  |
| *AtCPK28* | -2.67624 | 0.001017 |
| *AtCPK4* | -1.8105 | 0.032318 |
| *AtCPK6* | -1.77207 | 0.005785 |
| *AtCPK1* | -1.58869 | 0.007345 |
| *AtCPK2* | -1.56419 | 0.017867 |
| GNCG |  |  |
| *CNGC19* | -5.76803 | 0.001672 |
| Rboh |  |  |
| *RbohC* | -6.19656 | 0.02151 |
| *RbohD* | -2.86904 | 0.002968 |
| *RbohF* | -1.47496 | 0.041597 |
| MAPKKK |  |  |
| *MEKK1* | -1.95731 | 0.001834 |
| MAPKK |  |  |
| *MKK4* | -2.08301 | 0.003331 |
| *MKK5* | -1.55409 | 0.004236 |
| MAPK |  |  |
| *MPK11* | -3.68019 | 0.001659 |
| *MPK3* | -1.49632 | 0.022141 |
| *MPK6* | -1.06056 | 0.011904 |
| PROPEP |  |  |
| *PROPEP2* | -6.18334 | 0.001698 |
| *PROPEP3* | -5.5927 | 0.002962 |
| SA/PIP (Phytohormone biosynthesis pathway) |  |  |
| *EDS5* | -5.44339 | 0.000782 |
| *PBS3* | -4.88858 | 0.002298 |
| *ICS1* | -4.74602 | 0.001475 |
| Photosynthetic pathway genes |  |  |
| *Lhcb2* | 5.644727 | 0.000682 |
| *ATAF1* | 4.265528 | 0.002733 |
| *GUN4* | 4.175522 | 0.001681 |
| *Lhcb4* | 3.616073 | 0.000771 |

**Extended Data Fig. 5.**
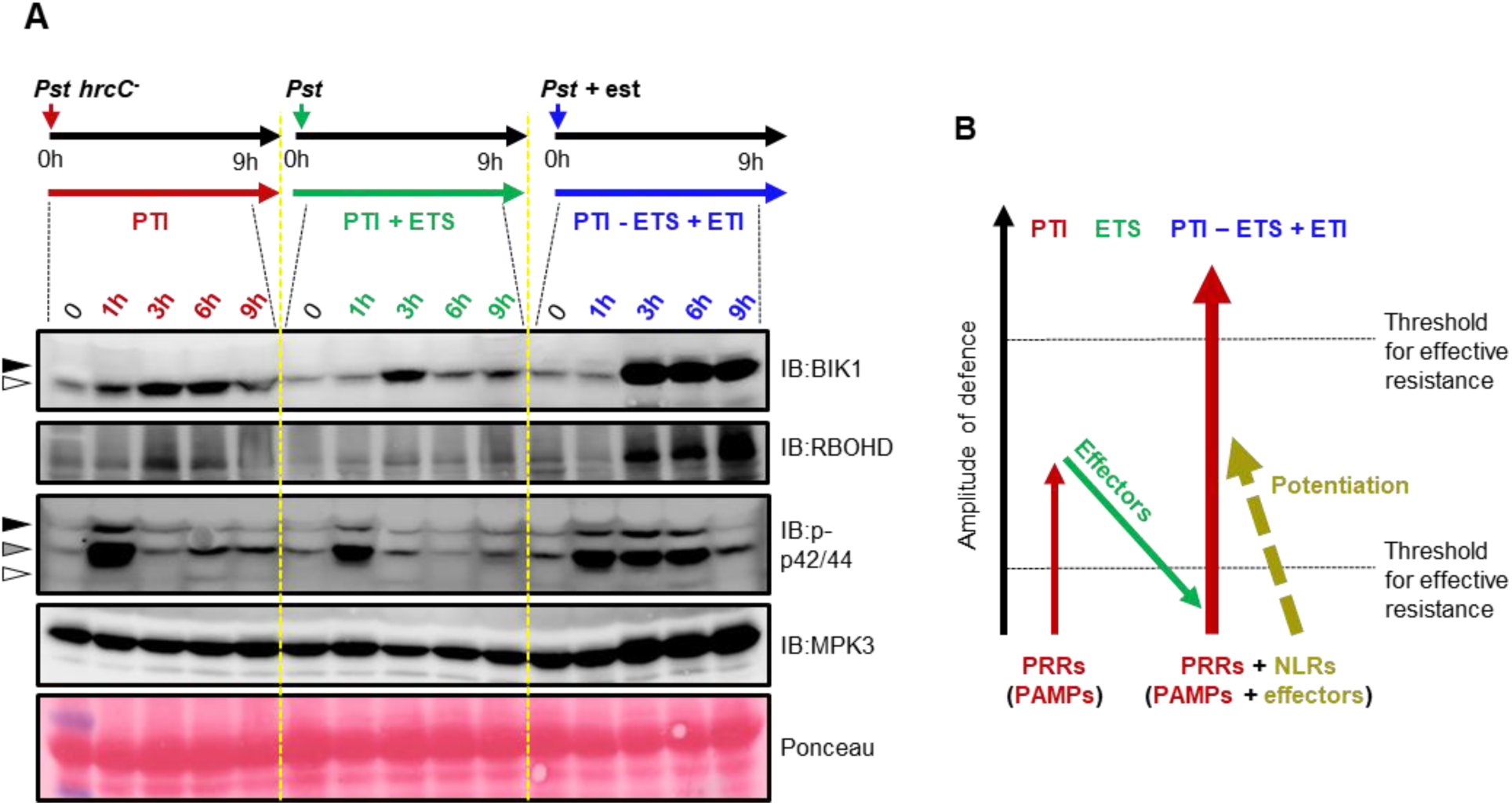
ETI ‘restores’ PTI from ETS. (A) 5-week old leaves of est:AvrRps4 were infiltrated with *Pst hrcC*^*-*^ (PTI), *Pst* (PTI + ETS), or 50 μM est *+ Pst hrcC*^*-*^ (PTI - ETS + ETI^AvrRps4^), and samples were collected at the indicated time points for protein extraction and immunoblotting. PTI leads to activation of MAPKs and accumulation of BIK1 and RbohD. *Pst* secretes effectors to block PTI (green). When PTI is coactivated with ETI, there is a stronger accumulation of MPK3, BIK1 and RbohD compared to that of PTI (blue). MAPKs activation by PTI is prolonged during PTI + ETI^AvrRps4^ through potentiation. Arrows in IB:BIK1 indicate the phosphorylation of BIK1 (black: pBIK1, white: BIK1). Arrows in IB:p-p42/44 indicate the corresponding MAP kinases (black: pMPK6, grey: pMPK3, white: pMPK4/11). Ponceau staining were used as loading control. (B) Updated version of the “zig-zag-zig” model. PAMPs from pathogen is detected by PRRs to trigger PTI. Successful pathogens secrete effectors to interfere with PTI, which leads to ETS. ETI is activated when effectors are recognized by NLRs. ETI restores and enhances PTI, which leads to effective resistance against virulent pathogens.

**Extended Data Fig. 6.**
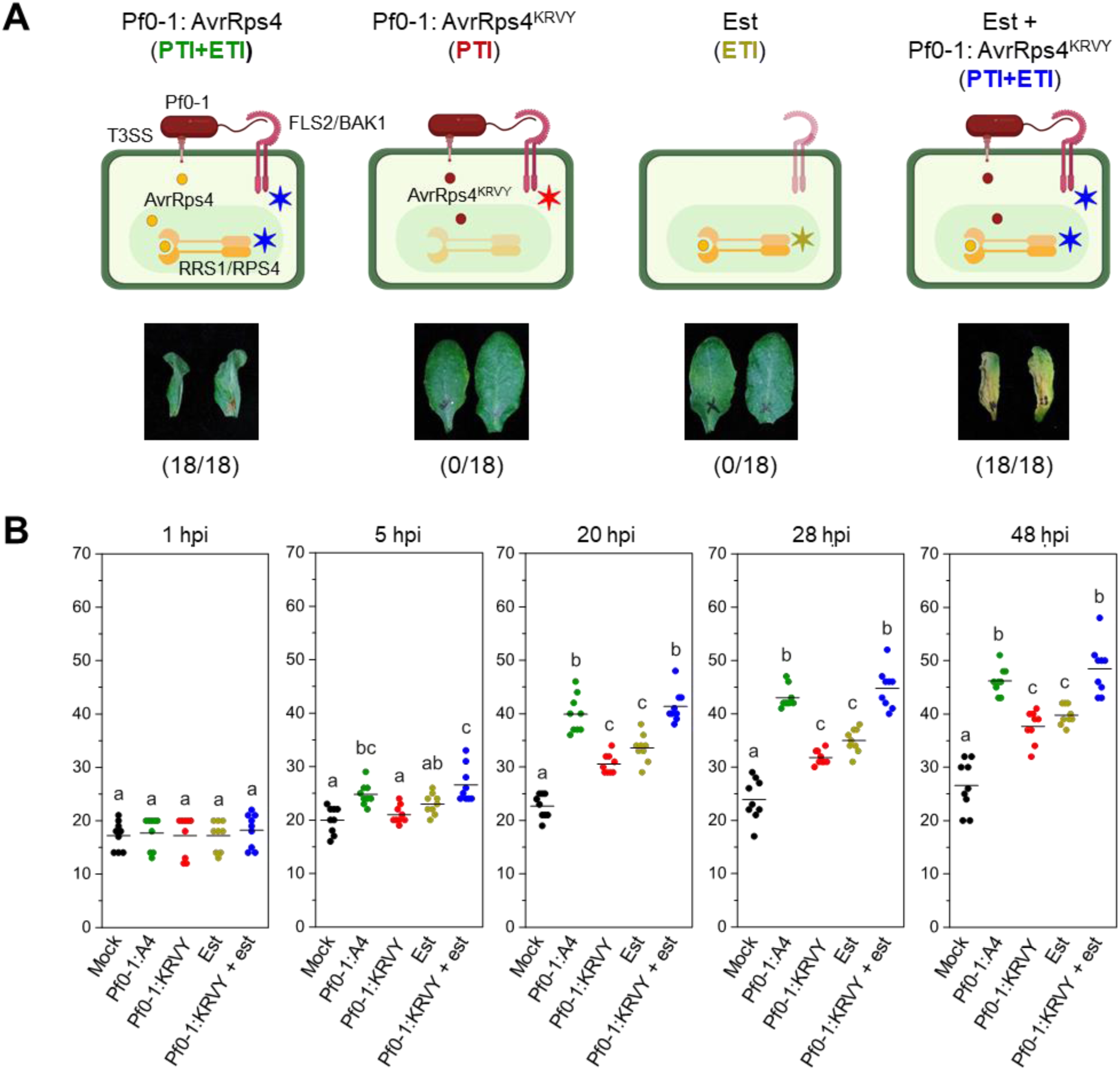
Potentiation of ETI^AvrRps4^-induced HR by PTI. (A) Pf0-1:AvrRps4 leads to macroscopic HR in est:AvrRps4 leaves. Both PTI (Pf0-1:AvrRps4^KRVY^) or ETI^AvrRps4^ (50 μM est) does not lead to macroscopic HR. Coactivation of PTI and ETI^AvrRps4^ (est + Pf0-1:AvrRps4^KRVY^) leads to macroscopic HR. The numbers indicate number of leaves displaying HR of the total number of leaves infiltrated. **✶** in the schematic diagrams indicate activated immune system. (B) Est:AvrRps4 leaves were hand-infiltrated with different solutions and electrolyte leakage was measured within 48 hpi. Combination of “PTI + ETI^AvrRps4^” (blue dots, est + Pf0-1:AvrRps4^KRVY^) leads to stronger electrolyte leakage compared to ETI^AvrRps4^ (est) or PTI (Pf0-1:AvrRps4^KRVY^) alone. Pf0-1:AvrRps4 (A4) acts as a positive control. Data points from 3 biological replicates were analyzed with one-way ANOVA followed by Tukey’s HSD test. Data points with different letters indicate P<0.01.

**Extended Data Fig. 7.**
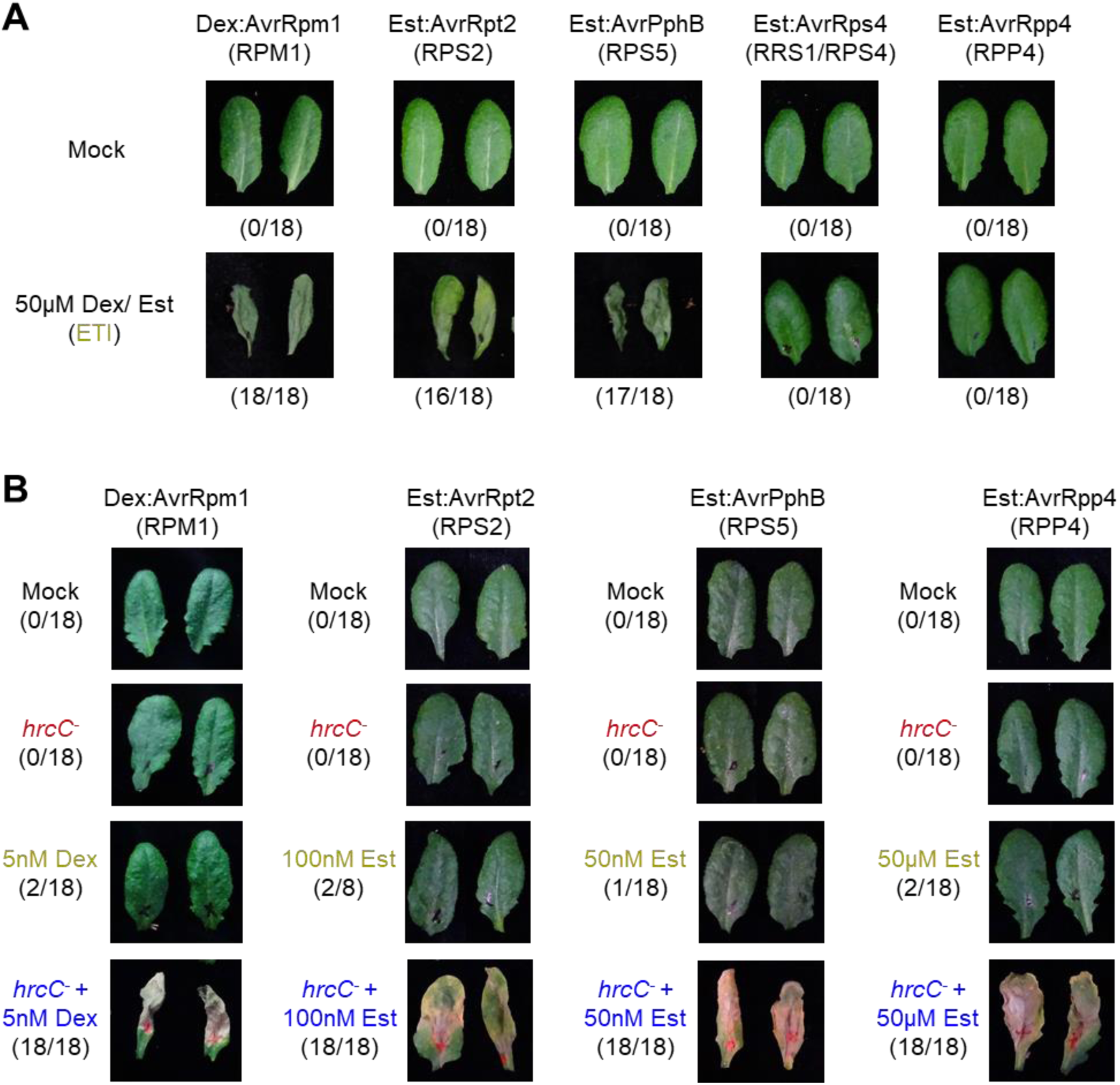
PTI potentiates HR induced by multiple NLRs. (A) 5-week old inducible-AvrRpm1 (dex:AvrRpm1, activates RPM1), AvrRpt2 (est:AvrRpt2, activates RPS2), AvrPphB (est:AvrPphB, activates RPS5), AvrRps4 (est:AvrRps4, activates RRS1/RPS4) and AvrRpp4 (est:AvrRpp4, activates RPP4) Arabidopsis leaves were infiltrated with 50 μM dex (for dex:AvrRpm1 only) or est. All pictures were taken 3 days post infiltration. The numbers indicate the number of leaves displaying HR of the total number of leaves infiltrated. (B) Combination of PTI+ETI leads to stronger macroscopic HR in inducible-AvrRpm1, AvrRpt2, AvrPphB and AvrRpp4 Arabidopsis lines. All pictures were taken 3 days post infiltration. The numbers indicate number of leaves displaying HR of the total number of leaves infiltrated.

**Extended Data Fig. 8.**
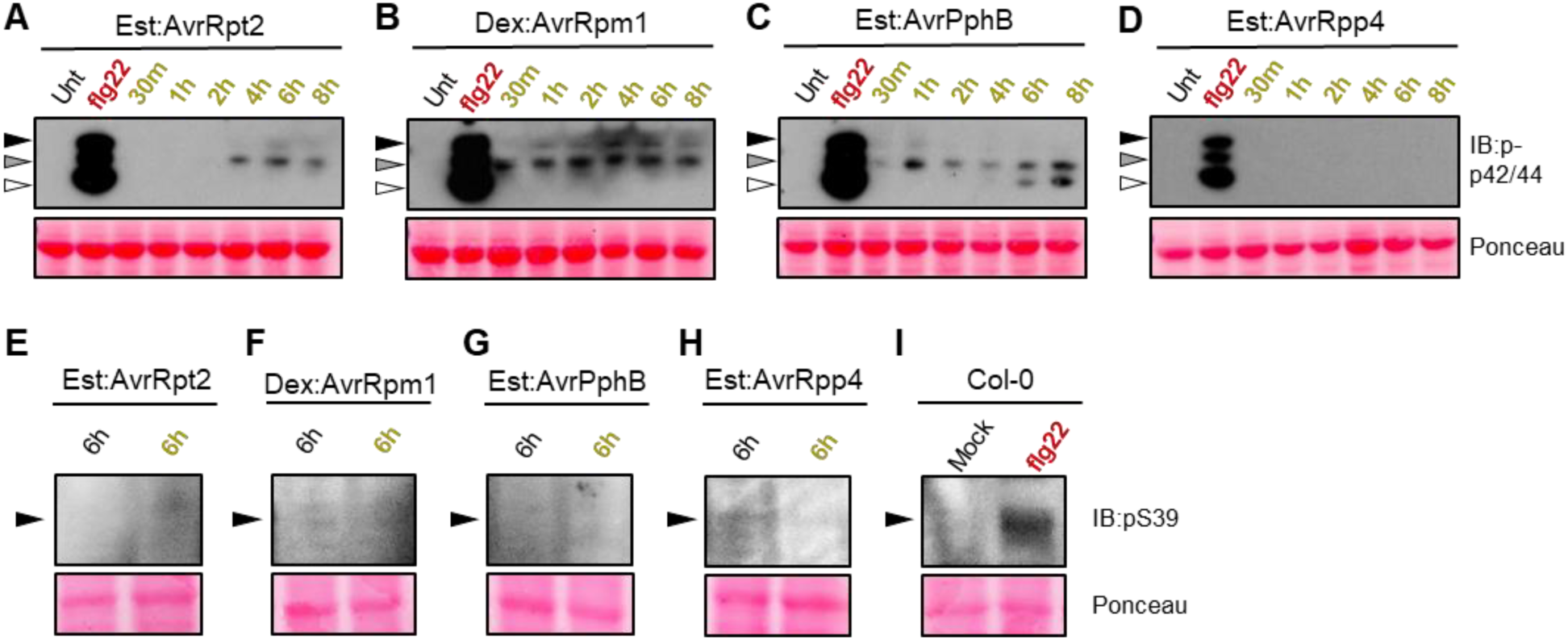
Phosphorylation of MAPKs and RbohD (S39) during ETI. Seedlings of (A) est:AvrRpt2, (B) dex:AvrRpm1, (C) est:AvrPphB and (D) est:AvrRpp4 lines were soaked in 50 μM dex (for dex:AvrRpm1) or 50 μM est, respectively, for indicated time points (dark yellow). Untreated (Unt) seedlings were used as negative control and seedlings treated with 100 nM flg22 for 15 minutes (red, flg22) were used as positive control. Arrowheads indicate the corresponding MAP kinases (black: pMPK6, grey: pMPK3, white: pMPK4/11). Seedlings of (E) est:AvrRpt2, (F) dex:AvrRpm1, (G) est:AvrPphB and (H) est:AvrRpp4 were soaked in mock solution (1% DMSO, black) or 50 μM dex or 50 μM est (dark yellow) for 6 h. (I) Col-0 seedlings treated with mock solution (1% DMSO) for 6 h (mock) were used as negative control and seedlings treated with 100 nM flg22 for 10 minutes (red, flg22) were used as positive control. For Extended Data Figures 1E and 8E-I, microsomal fraction from seedlings were isolated for immunoblotting. Ponceau staining were used as loading control for all the above experiments.

**Extended Data Fig. 9.**
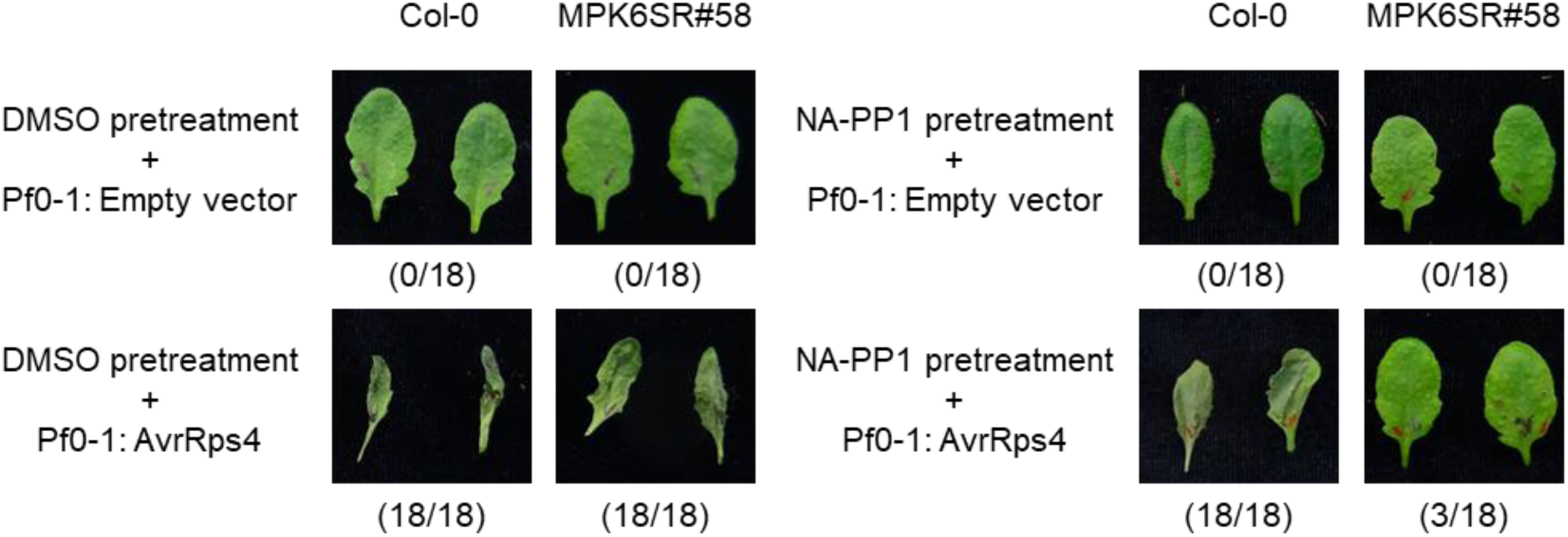
Inhibition of MPK6 in the MPK6SR#58 line attenuates HR induced by “PTI + ETI^AvrRps4^”. MPK6SR#58 (*mpk3 mpk6* P_MPK6_:MPK6^YG^) is a conditional *mpk3 mpk6* double mutant. MPK6^YG^ has a larger ATP binding pocket than MPK6^WT^ and is sensitive to the inhibitor 1-Naphthyl-PP1 (NA-PP1, ATP analog). Pre-treatment with NA-PP1 inhibits MPK6^YG^ and temporarily generates a *mpk3 mpk6* double mutant. Both Col-0 and MPK6SR#58 leaves were pre-infiltrated with either 1% DMSO (mock) or 1 μM NA-PP1. After 3 h, these leaves were infiltrated with either Pf0-1:Empty vector (triggers PTI) or Pf0-1:AvrRps4 (triggers “PTI + ETI^AvrRps4^”). With mock pre-treatment, Pf0-1:AvrRps4 infiltration leads to macroscopic HR in both Col-0 and MPKS6R#58. NA-PP1 pre-treatment attenuates HR caused by Pf0-1:AvrRps4 only in the MPK6SR#58 line. All pictures were taken one day post infiltration. The numbers indicate number of leaves displaying HR of the total number of leaves infiltrated.

## Notes

### Competing Interest Statement

The authors have declared no competing interest.

### Summary of Updates

This version of the manuscript has been revised to update the citations in the methods section.

